# Modelling the effect of migration on the localisation and spread of a gene drive

**DOI:** 10.1101/2023.04.02.535303

**Authors:** Benjamin Camm, Alexandre Fournier-Level

## Abstract

Gene drives have the potential to address pressing ecological issues. Through the super-Mendelian inheritance of a gene drive, a trait can be spread through a population even in spite of a fitness cost. This ability to spread is both its greatest quality and detractor. We may not want a gene drive to spread universally. If a gene drive were designed to cause the collapse of a pest population, it may inadvertently cause the collapse of the entire species. Migration is the mechanism through which a gene drive can spread to distant populations. Understanding its effect on the progression of a gene drive is crucial to our ability to control a gene drive. While migration can spread the gene drive to other populations, equally it can bring in other alleles to the population that may disrupt the progression of the gene drive. Through our deterministic migration gene drive model we can assess the conditions in which a gene drive is likely to spread to unintended populations, and if a gene drive is likely to be displaced by incoming alleles.

## Introduction

Gene drives have the potential to address pressing ecological issues through the function of super-Mendelian inheritance. A gene drive is able to spread a desired trait through a population of a species (Gantz & Bier, 2016). This may make a pest species more manageable, or eradicate it entirely (Legros et al., 2021; Prowse et al., 2017; Rode et al., 2019). A gene drive, by nature, is designed to spread, even in spite of a fitness cost (Gantz & Bier, 2016; Unckless et al., 2015). The invasive nature of a gene drive is both its greatest strength and biggest drawback. It needs to be considered that a gene drive designed to eradicate a pest species in one part of the world, may, inadvertently, eradicate the species in its native range (Dhole et al., 2019; Marshall & Hay, 2012; Sudweeks et al., 2019; Willis & Burt, 2021). There may be cases to be argued that some species could be driven to extinction, *Anopheles gambiae* comes to mind (Hammond et al., 2016), but many invasive pests in one part of the world are keystone species in another. It will be detrimental if a gene drive designed to control a pest population ends up causing global extinction of the species. Hence we need to have ways of reigning in the spread of a gene drive, and knowing when a gene drive is likely to spread through the effects of migration (Greenbaum et al., 2021).

While migration can assist in the spread of a gene drive (Dhole et al., 2018), it can also introduce resistant alleles into a population that can displace the gene drive. In the case of CRISPR/Cas gene drives where the homing efficiency is heavily influenced by the similarity between the gRNA sequence and target site (Champer et al., 2018; Hsu et al., 2013), migration of alleles with different target site sequences may introduce a degree of homing resistance into the population that will reduce the efficacy of the gene drive. If a sufficient amount of resistance alleles migrate into the population where a gene drive is trying to spread the gene drive may be displaced. These two aspects of migration, moving the gene drive out and moving other alleles in, are pivotal in establishing gene drive localisation. Gene drive localisation is an ideal goal for preliminary gene drive releases as it confines the effect of the gene drive spatially, which allows for long-term observation of the gene drive process (Akbari et al., 2013; Noble et al., 2019; Willis & Burt, 2021).

As gene drive technology is still in its infancy, there is not sufficient evidence to support the notion that a gene drive can be designed to be localised. Hence the burden falls to modelling to build confidence that a gene drive can be designed to be localised. Extensive modelling has been carried out in the gene drive field to better understand the processes that underlie the function of a gene drive (Champer et al., 2019; Drury et al., 2017; Rode et al., 2019; Unckless et al., 2015). These range from highly specific individual level simulations (Champer et al., 2019; Eckhoff et al., 2017; Wu et al., 2021) to more broadly applicable population genetics models (Drury et al., 2017; Rode et al., 2019; Unckless et al., 2015). Here we have used an established population genetics gene drive model (Drury et al., 2017) with the addition of migration to better understand how we can create gene drives that will remain localised.

An important aspect of gene drive modelling is that of equilibria. The equilibrium point of a gene drive is one where the frequency of the gene drive allele does not change (Unckless et al., 2015). These equilibria demarcate important design aspects as an equilibrium can be either stable or unstable. A stable equilibrium is one where the frequency of the gene drive will converge with the equilibrium point, whether it be above or below it. An unstable equilibrium point is one where the frequency of the gene drive will diverge away from it if it is above or below the point (Unckless et al., 2015). These equilibrium points are key to creating a localised gene drive. If we can keep the frequency of the gene drive below the unstable equilibrium in off-target populations then it should not fixate in the population. And conversely, if the frequency of the gene drive is able to get above the unstable equilibrium in the target population then the gene drive will fixate.

We have explored the effect of migration on both gene drive invasiveness and persistence in conjunction with selection, dominance, homing, and inbreeding under a mathematical deterministic framework. This system allows us to understand under what conditions and at what point a gene drive is likely to either spread, or be displaced due to the effects of migration.

## Methods

The basis for the pair of equations used in this modelling is from Drury *et al*. 2017 (corrected) with the resistance frequency equal to zero (Equation 1), which itself is derived from Unckless *et al*. 2015. We implement migration into the system in two ways: where gene drive alleles are migrating into the system (Figure 1A), and where wild-type alleles are migrating into the system (Figure 1B). The migration rate refers to what relative fraction of the receiving population is added. For simplicity, only one type of allele migrates into a population. We make this assumption with the understanding that either the gene drive has fixed in a population and is subsequently migrating to other populations. Or that other populations without the gene drive present are migrating into a population where the gene drive is trying to increase in frequency. Implementing migration in this way allows for a very flexible measure of migration. If we assume the populations are of equal size, then whatever fraction of the population leaves will be the same fraction that enters the population, which is then summed to one. If the populations are of different sizes, then the incoming fraction is adjusted by the relative sizes of the populations. It can even accommodate migration rates larger than one, where the incoming migrants represent a larger population than the receiving population. This is beneficial when considering changing population sizes. A gene drive could be used to suppress a population, thereby increasing the effect of migration into the population.

**Figure 1.**
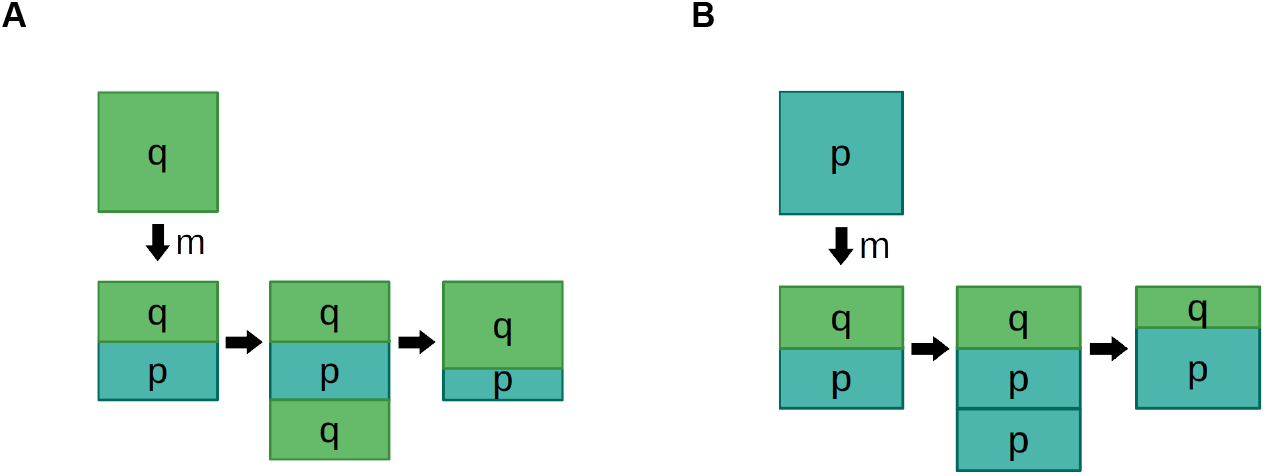
Examples of migration. A) Migration of a q allele into another population. B) Migration of a p allele into another population

When wild-type is migrating into a gene drive population, the relative frequency of the gene drive allele decreases by the factor 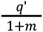 giving us Equation 2. When the gene drive is migrating into a wild-type population, the relative frequency of the gene drive allele increases by the facto r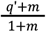 giving us Equation 3. Solving both of these equations for *q*_*m*_′ = *q*′ gives us two solutions, one for the stable and one for the unstable equilibria values, for WT into GD (Equation 4) and GD into WT (Equation 5). We ignore the trivial equilibria of frequency of the gene drive in both populations being one or zero.

To explore the sets of equations, solutions for the equations were solved over all combinations of a selection coefficient from 0.001 to 0.99 with a step of 0.001 and a migration rate of of 10 to the power of -5 to 2 with a step of 0.01. The conversion efficiency was either 0.05 or 0.3 for wild-type into gene drive or 0.05 or 0.5 for gene drive into wild-type and with either an inbreeding coefficient of 0.0 or 0.1.

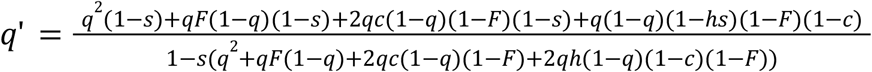

Equation 1. Gene drive model from Drury, with r=0

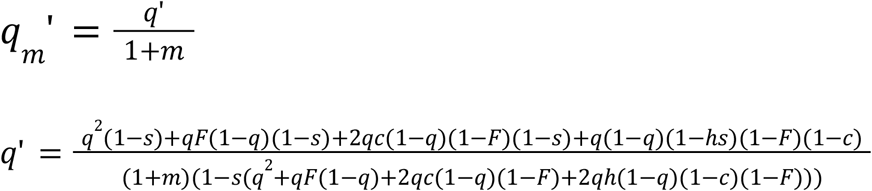

Equation 2. Equation modelling the migration of wild-types into a gene drive population.

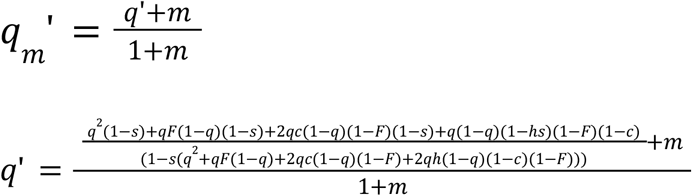

Equation 3. Equation modelling the migrations of gene drives into a wild-type population.

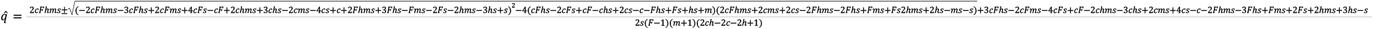

Equation 4. Equilibrium of wild-type into gene drive population.

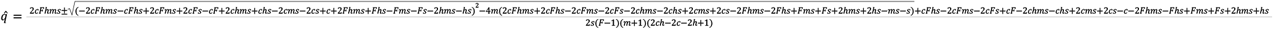

Equation 5. Equilibrium of gene drive into wild-type population.

## Results

### Stable and unstable equilibria

Our modelling demonstrated the effect migration has on a gene drive, with regard to both when a gene drive is likely to spread and when it will be displaced by resistance alleles. From solving the pair of equations (Equations 2 and 3) we saw that within the variable space, existed both a stable and unstable equilibrium. With the stable equilibrium being higher when the wild-type alleles are migrating into the gene drive population and lower when the gene drive alleles are migrating into the wild-type population (Figure 2). In both of these equations, we assume that the gene drive has already progressed and converted all the readily convertible alleles, with only resistance alleles remaining, hence the conversion efficiency is low.

**Figure 2.**
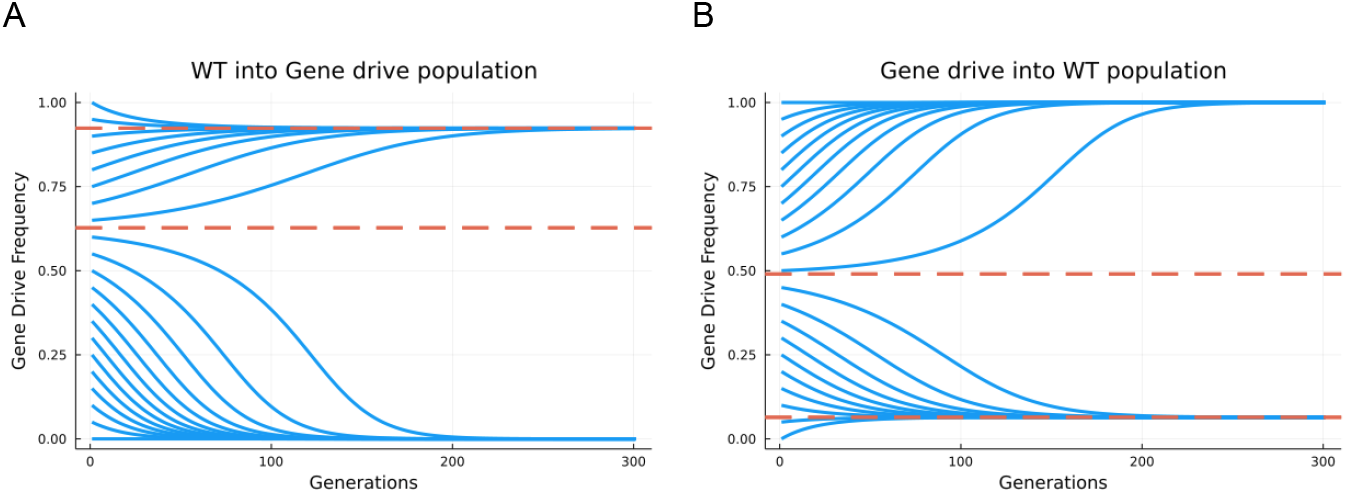
Equilibrium with migration. Each solid blue line represents a gene drive allele introduced into the population at different frequencies. Dashed orange lines represent equilibrium values. A) Modelling wild-type alleles moving into a gene drive population. Equilibrium points at 0.92 and 0.63. B) Modelling gene drive alleles moving into a wild-type population. Equilibrium points at 0.49 and 0.064. Both models have a selection coefficient of 0.1, dominance of 1.0, conversion efficiency of 0.05, inbreeding of 0.0 and a migration rate of 10^−2.5^.

When considering the wild-type migrating into a gene drive population we saw that when migration was non-zero, the gene drive was unable to fixate (Figure 2A). This is because every generation wild-type alleles were migrating into the population after homing had occurred, hence they were always present. The unstable equilibrium of the system defined the point above which the gene drive increased in frequency and below which the gene drive decreased in frequency. As the stable equilibrium was higher than the unstable equilibrium in this system, when the gene drive was above the unstable equilibrium, it would increase in frequency up to the stable equilibrium. When the frequency of the gene drive was below the unstable equilibrium, the gene drive decreased in frequency to zero. We saw a similar but reversed position of equilibria when considering the gene drive migrating into a wild-type population (Figure 2B). Here, the gene drive is never lost as migration at each generation of gene drive alleles maintains its presence in the population. Here the unstable equilibrium is higher than the stable equilibrium. We saw that when the frequency of the gene was above the unstable equilibrium it reached fixation. However, when the frequency of the gene drive was below the unstable equilibrium, it decreased until it reached the stable equilibrium point.

Now that we had an understanding of how the pair of equilibria affected the progression of the gene drive, with different styles of migration, we wanted to understand how the variables of the equation affected these equilibria. By solving Equations 2 and 3, resulting in Equations 4 and 5, and testing a range of each variable we saw how migration, selection coefficient, conversion efficiency and inbreeding affected these equilibria.

### Wild-type migrating into gene drive population

Firstly, considering the movement of wild-type alleles migrating into a gene drive population, we are investigating when a gene drive population is likely to resist incoming alleles. We saw that equilibria were only found when the migration rate was low, less than ∼10^−2.5^ and across the entire range of selection coefficients when the conversion efficiency was 0.05 (Figure 3A). When equilibria were not found (white space) it denoted conditions where the gene drive is lost. The unstable equilibrium values were regulated by the selection coefficient, whereby as the selection coefficient decreased, the equilibrium value decreased. Which means that with a lower selection coefficient, the gene drive needs to be above a lesser frequency in order to reach the stable equilibrium. The stable equilibrium was less affected by changes in selection coefficient. It was more affected by the migration rate of the simulation. With higher migration rates the stable equilibrium value decreased up to the point where the gene drive would be lost. When the conversion efficiency was increased from 0.05 to 0.30, there was an expansion in the range of valid equilibria (Figure 3B). The general trends in the equilibria remained the same, with the selection coefficient affecting the unstable equilibrium and migration rate affecting the stable equilibrium. However, with an increased conversion efficiency, the tolerance of the gene drive to higher selection coefficients and migration rates increased. Where previously a conversion efficiency of 0.05, selection coefficient of 0.4 and migration rate of 10^−3^ had no valid equilibrium meaning the gene drive would be lost to the wild-type. However, increasing the conversion efficiency to 0.3 creates an unstable equilibrium point at 0.554 and stable equilibrium at 0.997. This means that if the frequency of the gene drive is above 0.554 in the population it will be able to resist the migrating wild-type alleles and fixate. Increasing the inbreeding value from 0.0 to 0.1 reduces the variable space where equilibria are viable along the selection coefficient axis (Figure 3C). In these models the degree of dominance is set to 1, meaning that inbreeding increasing the frequency of heterozygotes is not changing the severity of selection against the gene drive, just reducing the efficacy of homing by reducing the frequency of gene drive heterozygotes.

**Figure 3.**
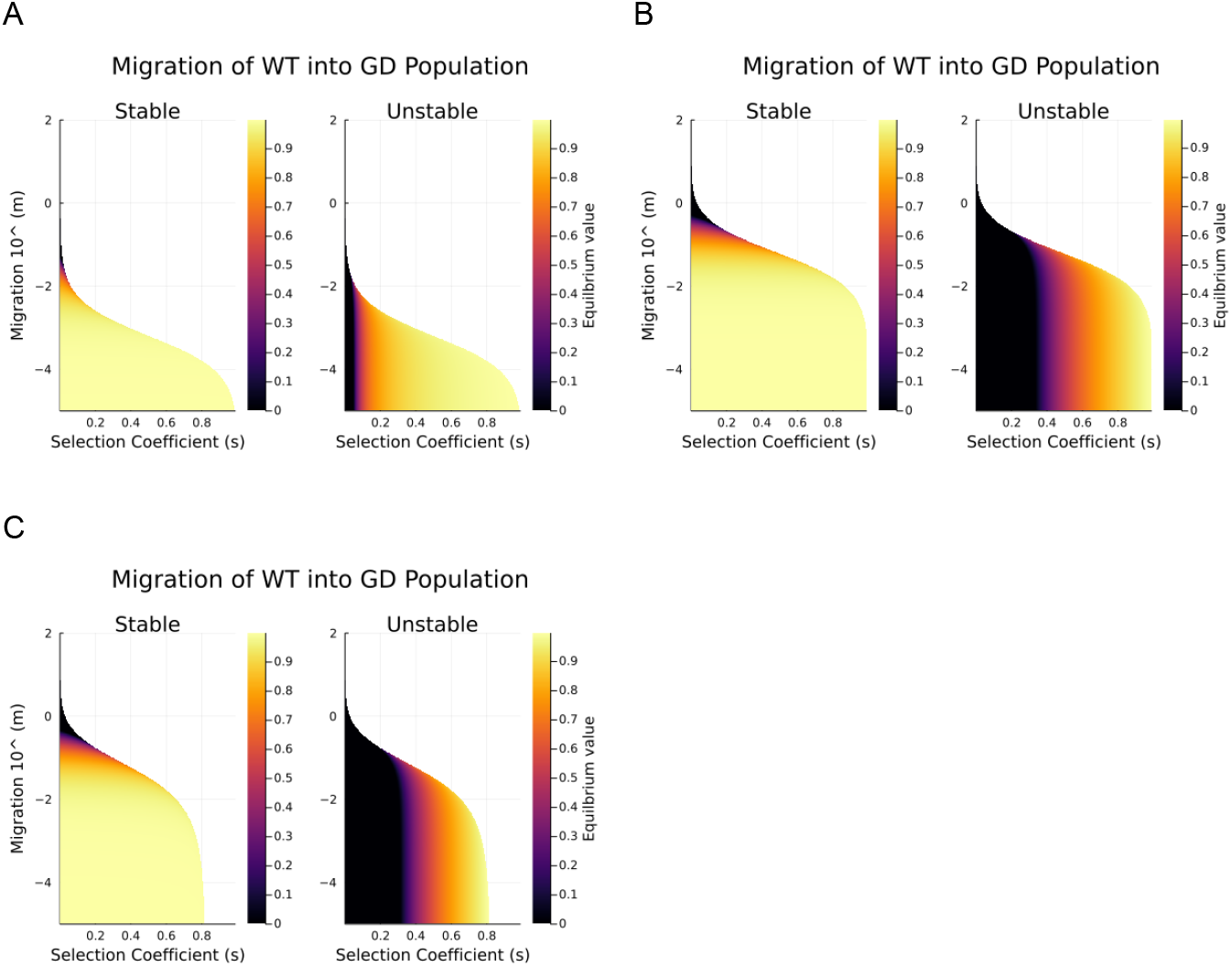
Stable and unstable equilibrium heatmaps for wild-type migrating into gene drive populations. White space is when no equilibrium is found, meaning that the gene drive allele is lost. A) Equilibrium points for a conversion efficiency of 0.05. B) Equilibrium points for a conversion efficiency of 0.3. C) Equilibrium points for a conversion efficiency of 0.3 and inbreeding of 0.1.

### Gene drive migrating into wild-type population

With regard to the migration of gene drive alleles into a wild-type population, we are trying to discern when a gene drive is likely to spread into unintended populations. Now that migration has changed from migrating wild-types to migrating gene drives, the equilibria have swapped positions, with the stable equilibrium below the unstable equilibrium. Now where no equilibria exist, it means that the gene drive has fixated. Again we saw how the selection coefficient and migration rate affected both the stable and unstable equilibria (Figure 4A). The unstable equilibrium is regulated by both the selection coefficient and the migration rate, as the change in equilibrium value runs along the edge of the heatmap. When the selection coefficient was less than ∼0.1, the stable equilibrium was 1.0 and the unstable equilibrium was 0.0. This meant that with such a low selection coefficient, regardless of the migration rate, the gene drive would fixate. When the conversion efficiency was increased to 0.5, we saw that the heat maps shifted along the selection coefficient axis (Figure 4B). The selection coefficient below which the gene drive would fixate regardless of the migration rate had increased to ∼0.4. Meaning the higher the conversion efficiency the more invasive a gene drive will be over a wider range of selection coefficients. The unstable equilibrium values decreased with a higher conversion efficiency, meaning that the frequency the gene drive needs to be at to fixate is less. The addition of inbreeding, from 0.0 to 0.1, erodes the equilibrium space but only for the unstable equilibrium (Figure 4C). The space eroded due to the addition of inbreeding denotes space where the gene drive is lost as the unstable equilibrium value is above 1 meaning the gene drive cannot be sustained.

**Figure 4.**
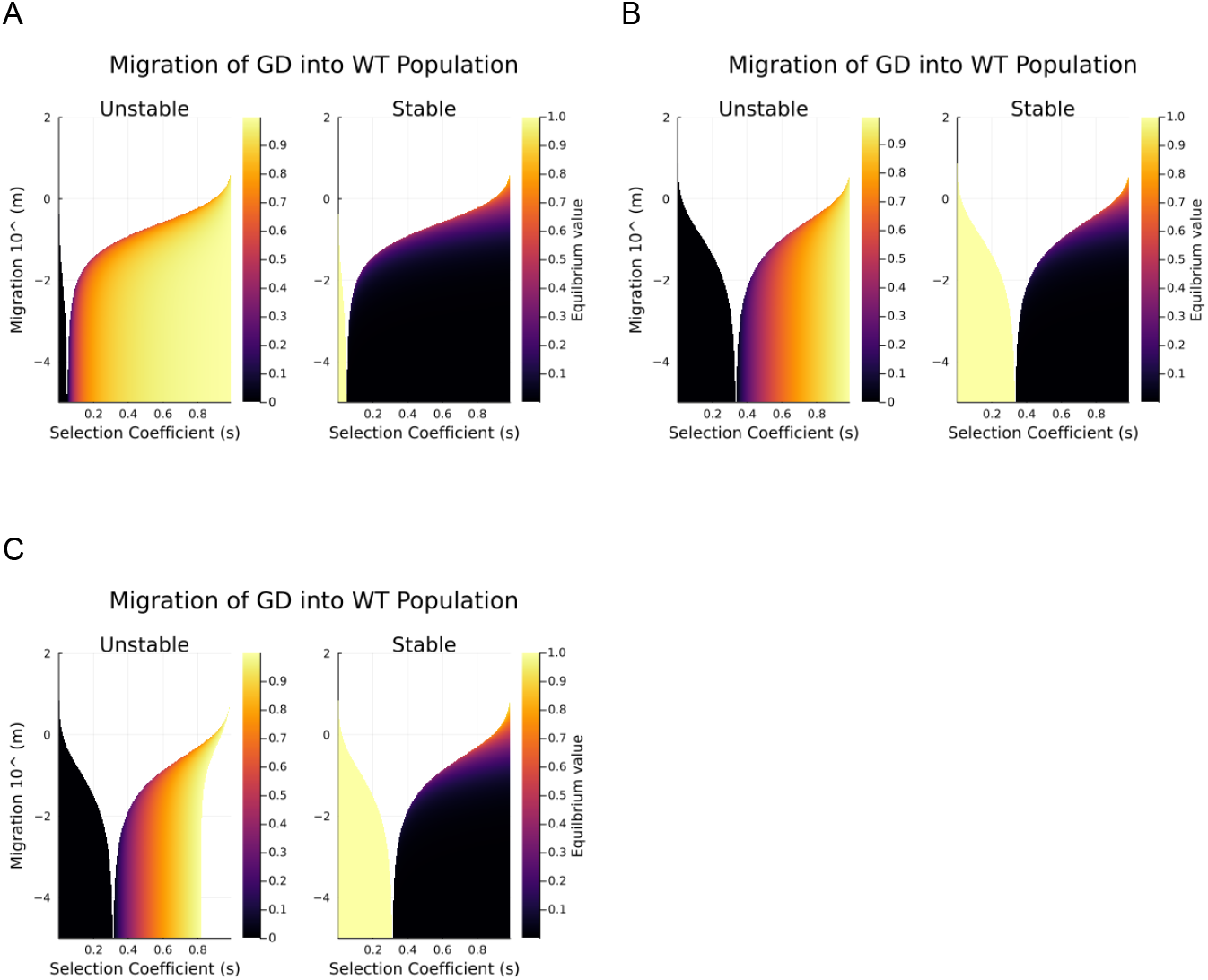
Stable and unstable equilibrium heatmaps for gene drive alleles migrating into wild-type populations. White space is when no equilibrium is found, meaning that the gene drive allele will fixate. A) Equilibrium points for a conversion efficiency of 0.05. B) Equilibrium points for a conversion efficiency of 0.5. C) Equilibrium points for a conversion efficiency of 0.5 and inbreeding of 0.1. The white space under the curve appearing from increasing inbreeding to 0.1 denotes when the gene drive is lost.

## Discussion

Gene drives are an invasive force that, while beneficial locally, may be detrimental globally (Marshall & Hay, 2012; Noble et al., 2019; Willis & Burt, 2021). As such, understanding the thresholds for a gene drive to spread through migration, or resist migration, are key to gene drive design and implementation (Greenbaum et al., 2021). Here we have explored a set of gene drive models to better understand how stable and unstable equilibria of a gene drive system are affected by the conversion efficiency, selection coefficient, dominance, inbreeding and most importantly, migration rate. Knowing these variables’ effect on the equilibria points will aid in creating gene drives that are less likely to spread beyond their boundaries, or gene drives that will be able to resist incoming resistance alleles.

Assumptions are made in this modelling framework for simplicity sake. With regard to the wild-type into a gene drive population, we only use a single conversion efficiency value, which we inevitably use as a proxy for the remaining resistance level of the population. We assume that a gene drive has spread in the population, converting all the alleles bar the resistance alleles which have a low conversion efficiency. And that the alleles from the migrating population are predominantly resistance alleles. Hence we model a low conversion efficiency. The same assumption is made for the gene drive into wild-type model. In this case we assume that the gene drive has spread in a population and reached near fixation and is now migrating into the wild-type population. In both these cases, the gene drive spread and converted all the readily convertible alleles, leaving only the resistant alleles behind.

The unstable equilibrium points of a gene drive are the tipping points of the system. Below which the frequency of the gene drive declines and above which the frequency of the gene drive increases. The stable equilibrium points of a gene drive are the coalescing points. A frequency around the stable equilibrium point will tend towards it. In previous gene drive modelling with only a single equilibrium point (Unckless et al., 2015) it was simpler to understand how a gene drive will interact with its equilibrium point. But when a gene drive system contains two different equilibria points it is less so. Depending on which equilibria, stable or unstable is higher, changes the dynamics of the system. As with the gene drive into the wild-type model, if the frequency of the gene drive is below the unstable equilibrium point it will converge to the stable equilibrium point, and if above it will fixate. However, in the wild-type into gene drive model, if the frequency of the gene drive is below the unstable equilibrium point then the gene drive will be lost, while if it is above the unstable equilibrium point it will converge to the stable equilibrium.

The wild-type into gene drive population modelling informs us when a gene drive is likely to be displaced due to migration. By knowing the unstable equilibrium point, for a given set of variables, we know that if the frequency of the gene drive was able to surpass this point then the gene drive will increase in frequency. We can then use the gene drive into a wild-type population model to tell us if the gene drive will spread into other populations, based on the variables of the model and the gene drive frequency in the other populations.

Safety remains a key concern in the implementation of gene drive technologies in the field of pest management (Akbari et al., 2015; Esvelt & Gemmell, 2017; Lunshof & Birnbaum, 2017). If we can abet fears of unintended spread through thorough modelling then it may assist in progressing the release of gene drives. Gene drives have such enormous potential for addressing ecological issues that plague our planet, the sooner we can safely implement gene drives the better we will be.

What needs to be considered with gene drive migration is how the population sizes may differ. A gene drive that causes population suppression will inadvertently increase the effect of migration into the suppressed population. Our modelling accommodates this as the migration rate is unbounded. Meaning it could support a migration rate of one, which is when the population receives a number of migrants equal to the current population size. This is relevant as when a population is suppressed, only a few migrants each generation will have a considerable effect on the genetic composition of the population. Hence the migration rate has a strong effect on the successful localisation or maintenance of a gene drive.

Our results here aim to better understand how migration will affect the stability of a gene drive. If the initial goals of gene drive implementation are to be localised, then accounting for the effect of migration is essential. We have shown here how the inclusion of migration changes a gene drive model from having a single equilibrium to having two alternate equilibria. Furthermore, we explored the effect each variable had on these equilibria as they are key to localising a gene drive. This research will aid in gene drive design and maximising the likelihood that a gene drive will act as intended.

## Notes

### Competing Interest Statement

The authors have declared no competing interest.

